# Genome-wide signals of drift and local adaptation during rapid lineage divergence in a songbird

**DOI:** 10.1101/243766

**Authors:** Guillermo Friis, Guillermo Fandos, Amanda Zellmer, John McCormack, Brant Faircloth, Borja Milá

**Affiliations:** National Museum of Natural Sciences, Spanish National Research Council (CSIC), Madrid 28006, Spain; Department of Biodiversity, Ecology and Evolution, Complutense University of Madrid, Madrid 28040, Spain; Department of Biology, Occidental College, Los Angeles, CA 90041, USA; Moore Laboratory of Zoology and Department of Biology, Occidental College, Los Angeles, CA 90041, USA; Department of Biological Sciences and Museum of Natural Science, Louisiana State University, Baton Rouge, LA 70803, USA

**Keywords:** local adaptation, isolation by adaptation, drift, selective gradients, redundancy analysis, postglacial expansion

## Abstract

The formation of independent evolutionary lineages involves neutral and selective factors, and understanding their relative roles in population divergence is a fundamental goal of speciation research. Correlations between allele frequencies and environmental variability can reveal the role of selection, yet the relative contribution of drift can be difficult to establish. Recently diversified systems such as that of the Oregon junco (Aves: Emberizidae) of western North America provide ideal scenarios to apply genetic-environment association analyses (GEA) while controlling for population structure. Genome-wide SNP loci analyses revealed marked genetic structure consisting of differentiated populations in isolated, dry southern mountain ranges, and more admixed recently expanded populations in humid northern latitudes. We used correlations between genomic and environmental variance to test for three specific modes of evolutionary divergence: (i) drift in geographic isolation, (ii) differentiation along continuous selective gradients, and (iii) isolation by adaptation. We found evidence of strong drift in southern mountains, but also signals of local adaptation in several populations, driven by temperature, precipitation, elevation and vegetation, especially when controlling for population history. We identified numerous variants under selection scattered across the genome, suggesting that local adaptation can promote rapid differentiation over short periods when acting over multiple independent loci.

## Introduction

Lineage diversification involves both selective and neutral factors, and elucidating their relative strengths and interactions in the process of evolutionary divergence is essential to understand the mechanisms underlying the early stages of speciation (Coyne and Orr 2004; Nosil 2012). Lineage differentiation can be driven by divergent natural selection (Darwin 1859; Coyne and Orr 2004), a mechanism that is at the core of ‘ecological speciation’ models, in which reproductive isolation arises as a by-product of cumulative, ecologically adaptive changes (Mayr 1947; Schluter 2000; Rundle and Nosil 2005). In turn, accumulation of genetic differences caused by drift in geographic isolation or in isolation-by-distance (IBD, Wright 1943; Wright 1946) has been proposed as a mode of divergence driven by neutral factors (Mayr 1954, 1963), a mechanism that can be particularly strong in populations of small effective size (e.g. Carson 1975; Templeton 1981; Uyeda et al. 2009). Selection and drift can also act jointly and even interact during evolutionary divergence in a number of ways. Geographic distance usually implies environmental differences that may drive adaptation to local conditions and ecological differentiation, even if populations are connected by moderate gene flow (Schluter 2000; Rundle and Nosil 2005). The evolution of local adaptation can in turn result in isolation-by-adaptation (IBA), a mode of divergence where adaptive changes lead to intrinsic barriers to gene flow, enabling genome-wide differentiation at both neutral and selected loci (Nosil et al. 2008; Funk et al. 2011). Consequently, geographic distance and ecological divergence may promote similar patterns of genetic diversity among populations, so that teasing apart the roles of neutral evolution and ecological adaptation in evolutionary diversification requires approaches that account for both environmental heterogeneity and neutral population structure (Wang and Bradburd 2014; Frichot et al. 2015; Rellstab et al. 2015; Forester et al. 2016).

Our capacity to assess the relative roles of adaptation and neutral differentiation in driving population divergence has benefitted from our increased ability to survey genome-wide variation thanks to the development of next generation sequencing (NGS) techniques (McCormack et al. 2013; Faria et al. 2014). The increasingly large number of loci afforded by NGS provides improved resolution to detect neutral population structure and patterns of gene flow among differentiated lineages. In addition, highly differentiated loci identified as outliers in an *F_ST_* distribution can be interpreted as potential targets of divergent selection standing out in a background of balanced or neutrally maintained genomic variation (Faria et al. 2014; Rellstab et al. 2015). However, methods of outlier detection relying solely on allele frequencies are sensitive to the confounding effects of historical factors, such as past sudden changes in population size or strong drift in small populations that may result in high rates of false positives (Edmonds et al. 2004; Kawecki and Ebert 2004; Billiard et al. 2005; Christmas et al. 2016). Moreover, changes in allele frequencies due to local adaptation can sometimes go undetected by outlier analyses (Pritchard and Di Rienzo 2010; Bierne et al. 2011; Rellstab et al. 2015; Forester et al. 2017). Alternative approaches that integrate environmental parameters by identifying allele frequencies that correlate with ecological variability have proven useful to detect signals of adaptation, especially when selective forces are weak (Frichot et al. 2015; Rellstab et al. 2015; Forester et al. 2017). These methods, known as genetic-environment association (GEA, Hedrick et al. 1976; Mitton et al. 1977) analyses, have the potential to reveal genetic patterns of differentiation due to local adaptation while testing for the role of multiple, specific environmental variables as drivers of selection (Forester et al. 2017). Importantly, GEA methods can correct for population history by controlling for general patterns of neutral genomic variation (Rellstab et al. 2015; Forester et al. 2016), allowing us to separate the respective effects of drift and selection in generating and maintaining variability. GEA analyses have greatly benefitted from the development of high-throughput sequencing techniques, resulting in a number of studies focusing on the genomic variability associated with environmental parameters in groups as diverse as plants (Lasky et al. 2012; Jones et al. 2013; De Kort et al. 2014; Nadeau et al. 2016; Sork et al. 2016), fungus (Ojeda Alayon et al. 2017), wolves (Forester et al. 2017) and birds (Manthey and Moyle 2015; Safran et al. 2016; Szulkin et al. 2016; Termignoni-García et al. 2017).

Recently diversified systems provide an ideal scenario for studying the relative roles of selective and neutral factors in incipient divergence and speciation. Specifically, GEA methods are particularly suitable when the system under study (i) is composed of closely related populations, among which the signals of selection are still recent and detectable; (ii) includes broad geographic distributions encompassing heterogeneous habitats across ecological clines (*i.e*. selective gradients) but also spatially discontinuous habitats so that adaptive and neutral divergence can be assessed in different spatial settings; (iii) shows large variability in the degree of geographical isolation among populations, from extensive gene flow to total isolation; and (iv) presents low variability in secondary sexual traits so that differential sexual selection can be ruled out as a major driver of population divergence.

The Oregon junco complex (*Junco hyemalis oreganus*) of western North America provides a particularly well suited system to carry out genome-environment association analysis. The complex originated recently as part of the postglacial radiation of dark-eyed juncos across North America following a northward recolonization of the continent as ice sheets retreated after the last glacial maximum, c.a. 18,000 years ago (Milá et al. 2007; Friis et al. 2016; Milá et al. 2016). Among dark-eyed junco forms, the Oregon junco group presents the highest variability in terms of phenotype and ecological range, encompassing a broad latitudinal range from Baja California to Alaska (Fig. 1A). All forms of the Oregon junco share a characteristic dark hood, yet there is considerable population variation in plumage color, mainly of the hood, dorsum and flanks, and the complex has been traditionally divided into at least 7 subspecific forms (Dwight 1918; Miller 1941; Nolan et al. 2002), which include, from south to north: *townsendi*, from the San Pedro Mártir mountains in northern Baja California, Mexico; *pontilis*, distributed just north of *townsendi* in the Sierra Juárez mountains, also in Baja California, Mexico; *thurberi*, from the mountains of southern California and Sierra Nevada; *pinosus*, a coastal form from central California, predominantly distributed in the Santa Cruz mountains; *montanus*, distributed across the interior of Oregon, Washington and British Columbia; *shufeldti*, a more coastal form from Oregon and Washington; and *oreganus* from coastal British Columbia and southern Alaska (Miller 1941; Nolan et al. 2002; Fig. 1A, Fig. 2A, Table 1).

**Figure 1.**
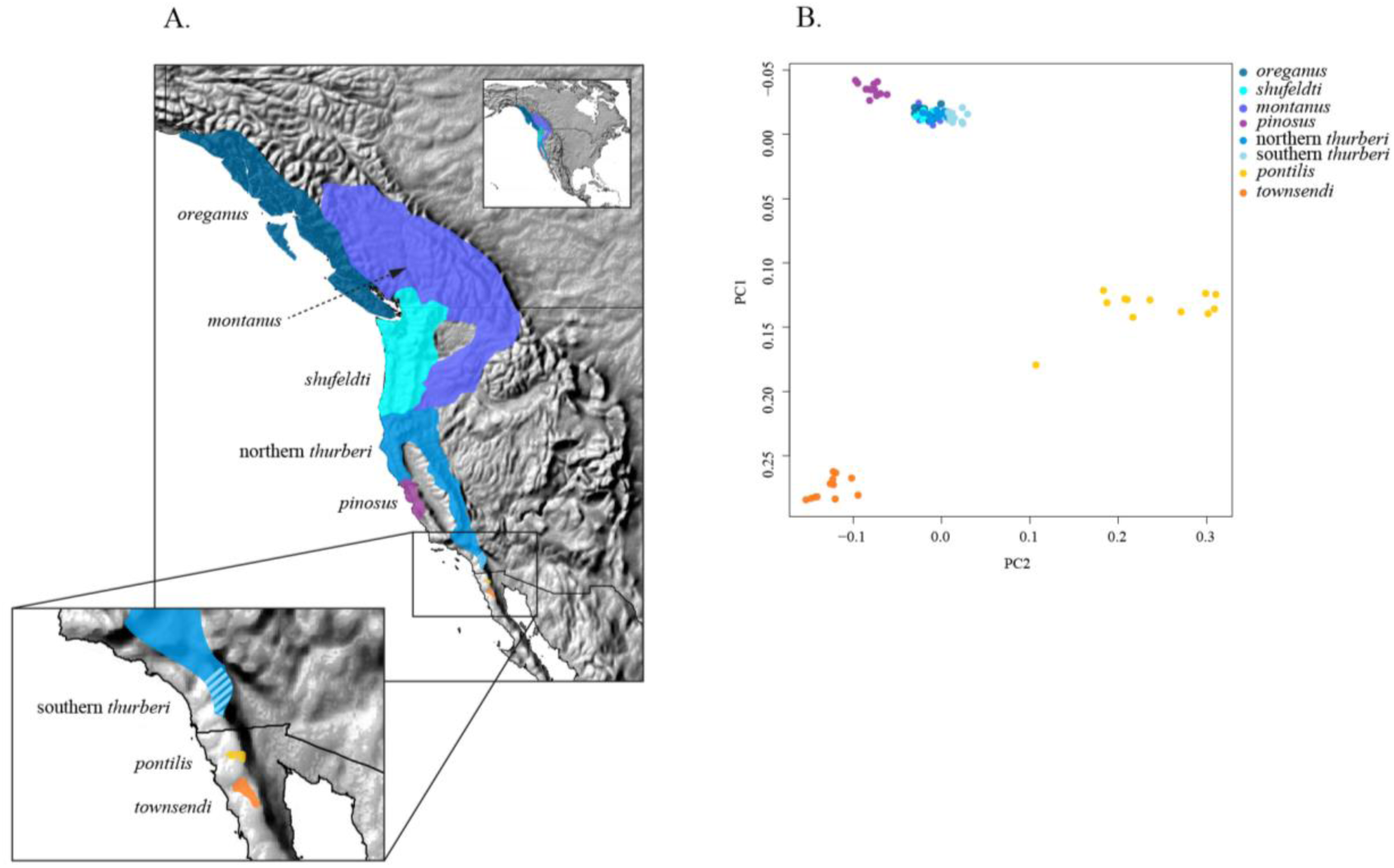
Geographic distribution and neutral genetic structure of the Oregon junco forms. (A) Breeding ranges of all the Oregon junco forms. (B) Genetic structure of Oregon junco forms based on a principal components analysis of independent, selectively neutral genome-wide SNPs.

**Figure 2.**
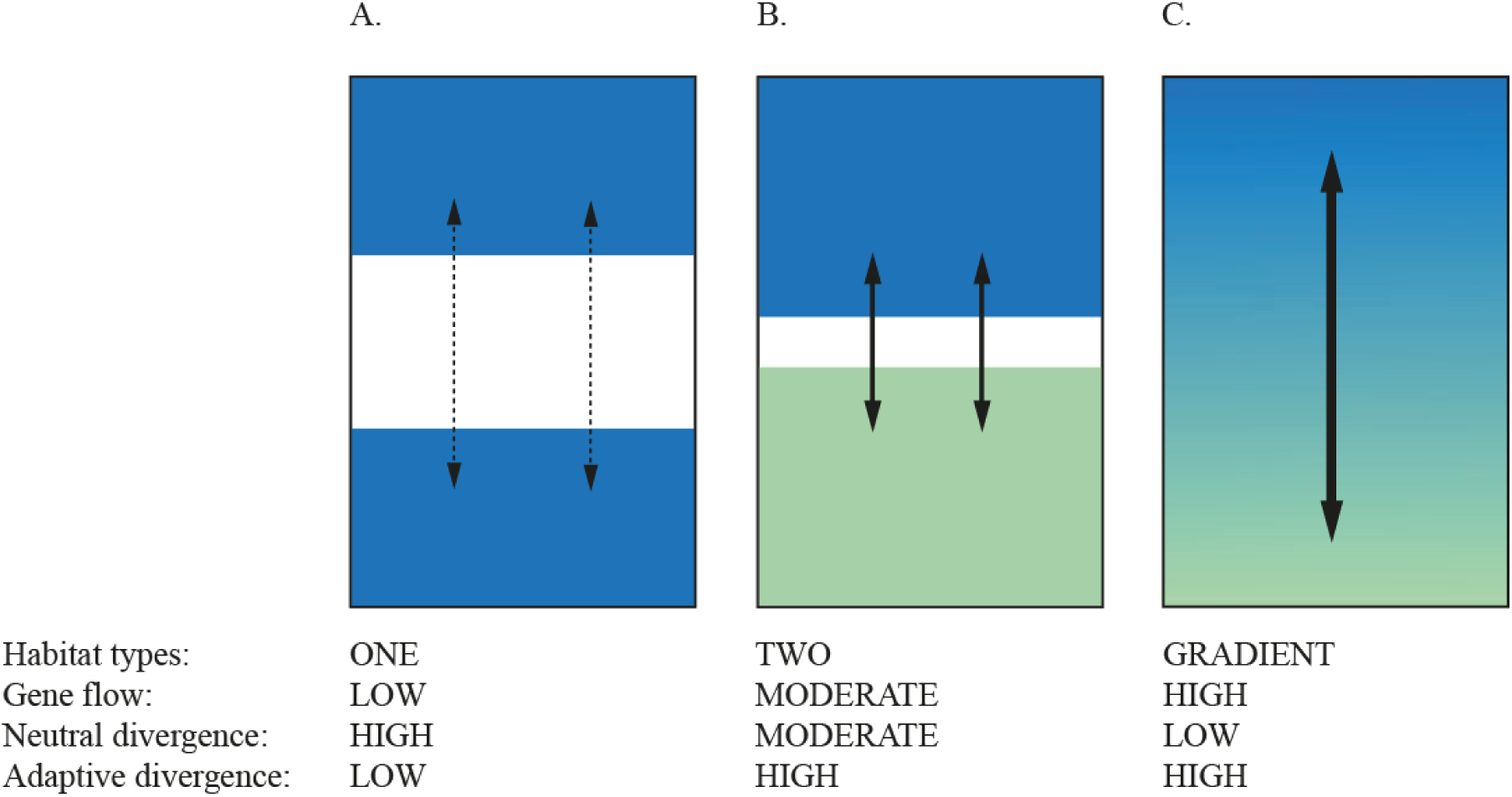
Expectations for neutral and adaptive divergence under different environmental and spatial configurations found across the Oregon junco distribution. (A) Geographically isolated populations in similar habitats. (B) Parapatric populations in ecologically divergent habitats. (C) Population continuum across a selective gradient.

**Figure 3.**
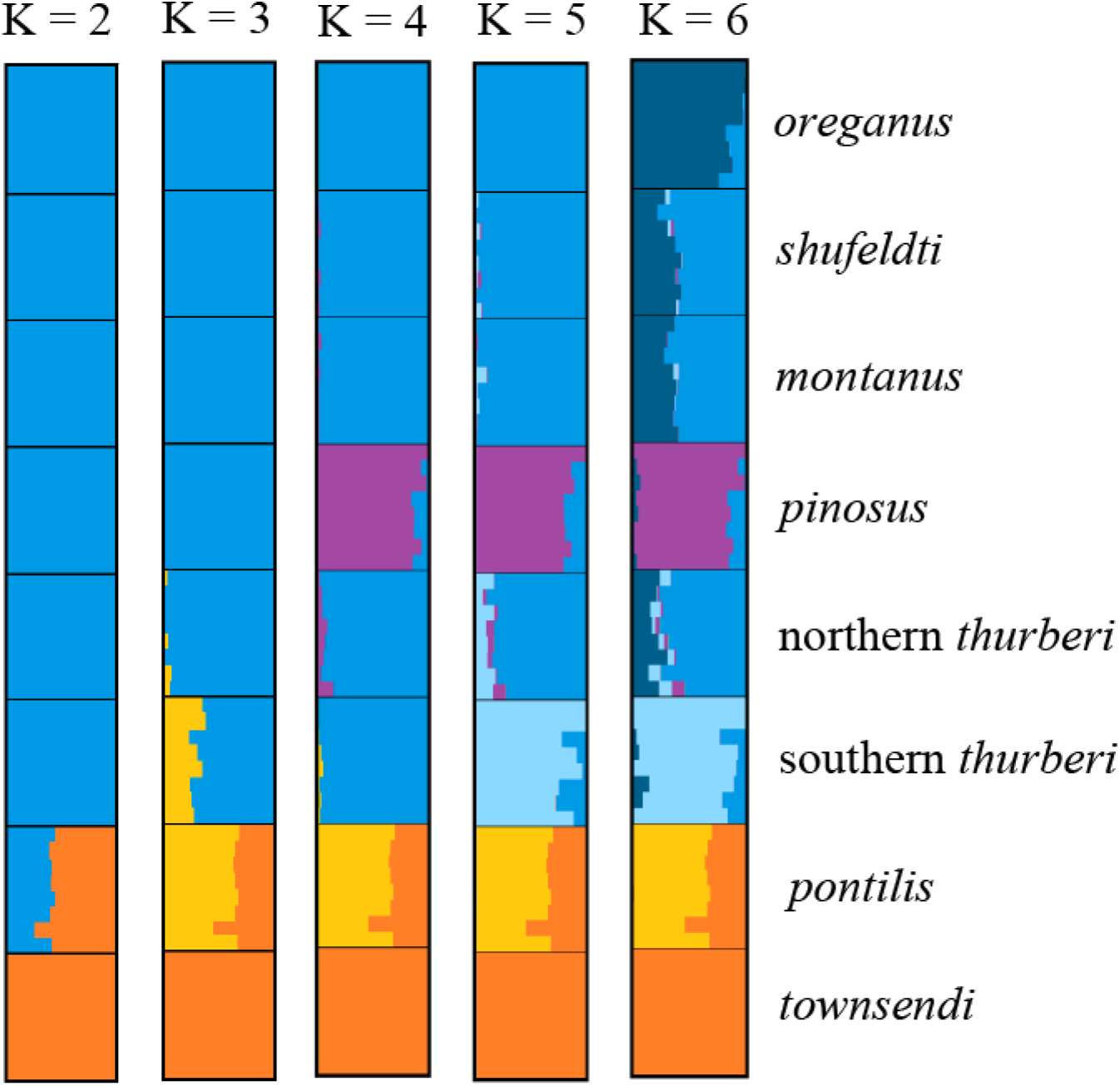
Genetic structure of the Oregon junco forms based on 34,367 selectively neutral genome-wide SNPs using the program STRUCTURE. Each horizontal bar corresponds to an individual, with colors corresponding to posterior assignment probabilities to each of a number of genetic clusters (K). Colors correspond approximately to those in Fig. 2A.

**Table 1.** Oregon junco forms and number of genotyped individuals per locality. State abbreviations are the following: British Columbia (BC) in Canada; Oregon (OR), California (CA) in the USA; Baja California Norte (BCN) in Mexico.

| *Form* | *State* | *Localities* | *Sequenced* |
| --- | --- | --- | --- |
| *oreganus* | BC | Banks Island, Porcher Island, Susan Island | 16 |
| *shufeldti* | OR | Willamette N.F. | 12 |
| *montanus* | OR | Wallowa N.F. | 16 |
| *northern thurberi* | CA | Tahoe | 18 |
| *southern thurberi* | CA | Mount Laguna | 17 |
| *pinosus* | CA | Santa Cruz Mountains | 16 |
| *pontilis* | BCN | Sierra Juárez | 16 |
| *townsendi* | BCN | Sierra San Pedro Mártir | 16 |
| Total |  |  | 127 |

The diverse spatial configuration of populations and environmental variability across the Oregon junco distribution are critical aspects that will affect our capacity to disentangle the roles of adaptive and neutral factors in explaining genomic variance. Here we use a conceptual framework to classify into three main settings the distinct spatial scenarios observable in the Oregon junco with respect to gene flow and environmental variation. These include (i) geographically isolated populations in similar habitats, as in the case of the Baja California *townsendi* and *pontilis* forms, where low levels of local adaptation and low rates of gene flow should result in limited adaptive divergence and high neutral divergence by drift (Fig. 2A); (ii) parapatric populations under divergent ecological conditions, as exemplified by the *pinosus* and *thurberi* forms in California, where divergence is expected to increase due to local adaptation, while geographic proximity and moderate gene flow should lead to intermediate levels of neutral differentiation by drift (Fig. 2B); and (iii) populations found along a continuous environmental gradient, as in the northernmost forms of Oregon junco (*thurberi*, *shufeldti*, *montanus* and *oreganus*) where neutral divergence is expected to be low due to high levels of gene flow, while local adaptation along the gradient may result in a pattern of high differentiation in adaptive variation (Fig. 2C).

Here we use the Oregon junco complex to study how geographic isolation, population history, and local ecological adaptation have driven population differentiation across the range, using extensive population sampling and genome-wide SNPs obtained from ‘genotyping by sequencing’ (GBS, Elshire et al. 2011). A draft consensus genome of junco has also been sequenced and assembled to be used as a reference. We first assess patterns of neutral genetic structure across the complex using selectively neutral SNPs, and we then look for correlations between environmental variables and allele frequencies across the Oregon junco distribution using redundancy analysis (RDA). We also use climatic variables to test for significant niche divergence while controlling for spatial autocorrelation. Finally, we map GBS sequences harboring significant outlier SNP loci to the zebra finch (*Taeniopygia guttata*) reference genome in order to recover the chromosomal position of polymorphic sites and explore how adaptive variation is distributed across the genome.

## Materials and methods

### Population sampling

Oregon junco populations were sampled across the geographical range of the species using mist nets in order to obtain biological samples for DNA extraction (Fig. 1A). Each captured individual was aged, sexed, and marked with a numbered aluminum band. A blood sample was collected by venipuncture of the sub-brachial vein and stored in Queen’s lysis buffer (Seutin 1991) or absolute ethanol at −80°C in the laboratory. All sampling activities were conducted in compliance with Animal Care and Use Program regulations at the University of California Los Angeles, and with state and federal scientific collecting permits in the USA and Mexico. A high-quality tissue sample for whole-genome sequencing was obtained from a red-backed junco (*J. hyemalis dorsalis*) specimen (MLZ 69090) provided by the Moore Laboratory of Zoology, Occidental College, that was collected at Dude Mountain, Coconino County, Arizona. Genomic DNA was extracted from blood and tissue samples using a Qiagen DNeasy kit (Qiagen™, Valencia, CA).

### Whole-genome sequencing and assembly

We assembled a draft dark-eyed junco genome by combining low-coverage genomes of eight different junco individuals, which we then used as a conspecific reference in the SNP calling process. Libraries for seven of the genomes were prepared with the Kapa Library Preparation Kit (Kapa Biosystems, Inc.) using TruSeq-style adapters (Faircloth and Glenn 2012). They were pooled after random shearing and individual barcoding and sequenced in a single lane of an Illumina HiSeq platform. The eighth genome was sequenced at a higher coverage by means of two 101-bp paired-end shotgun libraries and two 101-bp mate-paired libraries with insert sizes of 8 Kb in length at Macrogen Inc. The TruSeq Nano DNA Kit (Illumina) was used for the preparation of the shotgun libraries, while the mate-paired libraries were prepared with Nextera Mate Pair Kit (Illumina). We used FASTQC (Andrews 2010) to evaluate the quality of the sequenced data, and quality filtering was carried out with NextClip (Leggett et al. 2013) in the case of the mate paired libraries. For the rest of them we used Trimmomatic (Bolger et al. 2014), applying a sliding window filtering approach with a size of 4 bp and a phred quality score threshold of 25. We also set a minimum length of 50 bp, below which reads were filtered out after trimming. We used the software SOAPDENOVO2 (Luo et al. 2012) to perform the assembly. The average insert size for each library was estimated in a preliminary run, and we set a Kmer size of 27 and minimum edge coverage of 2. Gaps that emerged during the scaffolding process were removed with the GapCloser tool from SOAPDENOVO2. Finally, we filtered out all the scaffolds shorter than 500 bp so the genome was functional as a mapping reference. The final assembled genome had 37,904 scaffolds with an N50 of 147,816 bases, a L50 of 1,951 scaffolds, a total size of 1.09 Gb, 17.5 Mb of missing sites, and an overall coverage of ~56X, as computed with VCFTOOLS version 0.1.13 (Danecek et al. 2011).

### Genotyping-by-sequencing

We used genotyping-by-sequencing (Elshire *et al*. 2011) to obtain individual genotypes from 127 Oregon juncos belonging to the following subspecific taxa: *townsendi* (n=16), *pontilis* (n=16), *thurberi* (n=35), *pinosus* (n=16), *montanus* (n=16), *shufeldti* (n=12), *oreganus* (n=16) (Table 1). GBS libraries were prepared and sequenced at Cornell University’s Institute for Genomic Diversity, using the restriction enzyme PstI for digestion. Sequencing of the 127 individually-barcoded libraries was carried out in three different lanes (along with other 149 junco samples intended for other studies) of an Illumina HiSeq 2000, resulting in an average of 229.2 million good single-end reads 100 bp in length per lane. Samples of the same form were distributed among at least two of the lanes to avoid sequencing bias, except in the case of *shufeldti* individuals, which were all introduced in the last set of sequenced samples.

### Alignment and variant calling

We evaluated GBS read quality using FASTQC after sorting them by individual with FASTXTOOLKIT (Gordon and Hannon 2010), and performed the trimming and filtering treatment using PRINSEQ (Schmieder and Edwards 2011). All resulting reads were 69 bp long and had a mean genotyping phred quality score of at least 30, with no positions below 20. The reads were then mapped against the assembled junco genome using the mem algorithm in the Burrows-Wheeler Aligner (BWA, Li and Durbin 2009). Two samples were excluded at this step due to too low read mapping (1 *thurberi* and 1 *montanus*). We used the Genome Analysis Toolkit (GATK, McKenna et al. 2010) version 3.6-0 to realign reads around indels using the IndelRealigner tool and then we applied the HaplotypeCaller tool to call the individual genotypes. We finally used the GenotypeGVCFs tool to gather all the per-sample GVCFs files generated in the previous step and produce a set of joint-called SNPs and indels in the variant call format (*vcf*) (GATK Best Practices, DePristo et al. 2011; Auwera et al. 2013). Because GBS data do not provide enough coverage for base quality score recalibration, we used VCFTOOLS to implement a ‘hard filtering’ process, customized for each of the downstream analyses (Table 2).

**Table 2.** SNP data matrices used in each analyses. General filters included a depth range from 2 to 100 and a p-value for Hardy-Weinberg deviation of 10^−4^.

| Analysis | Number of<br>samples | Min. phred<br>score | MAF<br>threshold | Number<br>of SNPs | Allowed<br>missing data |
| --- | --- | --- | --- | --- | --- |
| STRUCTURE | 64 | 70 | 0.02 | 34,367 | 10% |
| PCA | 88 | 40 | 0.02 | 9,436 | 0% |
| RDA: |  |  |  |  |  |
| On all *loci* | 88 | 40 | 0.02 | 11,261 | 0% |
| On BayeScan outliers | 88 | 40 | 0.02 | 87 | 0% |
| Genome Scans: |  |  |  |  |  |
| All lineages | 88 | 40 | 0.02 | 29,868 | 75% |
| *townsendi* vs. *pontilis* | 24 | 40 | 0.05 | 22,773 | 75% |
| *townsendi* vs. *thurberi* | 24 | 40 | 0.05 | 22,516 | 75% |

### Genetic structure analyses

To explore genome-wide population structure among all Oregon junco forms, we ran a principal components analysis (PCA) based on SNP data. Using VCFTOOLS we retained all samples with less than 25% missing data after a ‘soft filtering’ (coverage range between 2 and 100, minimum phred quality score of 40), resulting in a dataset of 88 samples, and between 8 and 12 individuals per population (24 in the case of *thurberi*). We filtered out all the sites with any non-genotyped individuals or a minor allele frequency (MAF) below 0.02. We also applied a threshold for SNPs showing highly significant deviations from Hardy-Weinberg equilibrium (HWE) with a p-value of 10^−4^ to filter out false variants arisen by the alignment of paralogous loci, resulting in a matrix of 11,261 variants. We then excluded SNPs putatively under selection using BayeScan (Foll and Gaggiotti 2008). BayeScan computes per-SNP *F_ST_* scores and decomposes them into a population-specific component shared by all loci that approximates population related effects, and a locus-specific component shared by all populations, which accounts for selection. The program compares two models of divergence, with and without selection, and assumes a departure from neutrality when the locus-specific component is necessary to explain a given diversity pattern (Foll 2012). We used BayeScan with default settings and a thinning interval size to 100 to ensure convergence. For each SNP we obtained the posterior probability for the selection model and the *F_ST_* coefficient averaged over populations. For outlier detection and exclusion, we implemented a false discovery rate of 0.3. To filter out the SNPs under linkage disequilibrium (LD) we used the function snpgdsLDpruning from the SNPrelate package (Zheng 2012) in R Studio (R Studio Team 2015) version 1.0.136 with R (R Core Team 2013) version 3.2.2. We applied the correlation coefficient method with a threshold of 0.2 (method =“corr”, ld.threshold=0.2), resulting in a final data matrix of 9,436 SNP loci (Table 2). We then used the function snpgdsPCA also available in SNPrelate to perform the PCA and obtain the eigenvectors to be plotted.

We examined population structure with STRUCTURE (Pritchard et al. 2000), using a smaller, more heavily filtered SNP data matrix to reduce the computational time of the analysis. Using VCFTOOLS, we retained the eight samples of each population (16 in the case of *thurberi*) with the lowest proportion of missing sites for a final number of 64 samples. We constructed a data matrix of biallelic SNPs excluding those out of a range of coverage between 2 and 100, or with a genotyping phred quality score below 70. Positions with less than 90% of individuals genotyped were removed from the data matrix, along with those presenting a MAF below 0.02. Once again, we implemented a threshold for SNPs showing highly significant deviations from Hardy-Weinberg equilibrium (HWE) with a p-value of 10^−4^, and performed the filtering for non-neutral positions and linkage disequilibrium exactly as done for the PCA, to obtain a final data matrix of 34,367 SNP loci (Table 2). We converted the *vcf* file to STRUCTURE format using PGDspider (Lischer and Excoffier 2012) version 2.0.5.1. Bash scripts to perform the analyses were created with STRAUTO (Chhatre and Emerson 2016) and we ran the program five times per K, with K ranging from 1 to 10 after running a preliminary analysis to infer the lambda value. The burn-in was set to 50K iterations and the analysis ran for an additional 100K iterations. Similarity scores among runs and graphics were computed with CLUMPAK (Kopelman et al. 2015).

### Redundancy analysis and variance partition

When applying GEA methods there are two main potentially confounding effects related to neutral factors: (i) structure among populations derived from strong drift in isolation may result in genetic patterns similar to those related to adaptive divergence; and (ii) demographic expansions along latitudinal axes may create gradients of allele frequencies at neutral loci correlated with latitude, that in turn would correlate with any environmental variable that changes with latitude, mimicking a pattern of selective sweep and local adaptation (Excoffier and Ray 2008; Excoffier et al. 2009; Rellstab et al. 2015; Forester et al. 2016). Redundancy analysis (Van Den Wollenberg 1977; Legendre and Legendre 1998) is a canonical ordination method that allows computing the variance of a set of response variables explained by a number of constraining or explanatory variables. In addition, partial RDA enables computation of this shared variance between two sets of variables while conditioning or holding constant the effects of a third set of covariables. Here we used RDA and partial RDA as implemented in the R package vegan (Oksanen et al. 2016) to explore the associations between genetic variability and environmental data. Ecological data were obtained from 7 of the 19 variables available in the BioClim database (Hijmans et al. 2005), specifically chosen in accordance to their relevance to junco ecology (Miller 1941; Nolan et al. 2002). They measured mean temperature and precipitation over the year (BIO1 and BIO12); mean temperature and precipitation over the warmest quarter (BIO10 and BIO18), which corresponds to the birds’ breeding season; isothermality, referring to how the range of day-to-night temperature differs from the range of summer-to-winter, where a value of 100 indicates equality between them; and seasonality of temperature and precipitation (BIO4 and BIO15). We also included three vegetation variables from the Moderate Resolution Imaging Spectroradiometer (MODIS) satellites as available in https://modis.gsfc.nasa.gov: percent tree cover (TREE), Normalized Difference Vegetation Index (NDVI, a measure of canopy greenness), and NDVI’s annual standard deviation (std_NDVI). Finally, we included the high-quality elevation data provided by the NASA Shuttle Radar Topographic Mission (ELEV), downloadable from http://www2.jpl.nasa.gov/srtm for a total of eleven variables (Table 3). All ecological variables were centered and standardized. Following Blanchet et al. (2008), we implemented a forward selection method using the forward.sel function from the R-package packfor (Dray et al. 2009) to reduce the number of variables in the model. This procedure applies two stopping criteria: a significance level for each tested variable, which we set at 0.01; and a maximum limit for global adjusted R^2^, equal to the adjusted R^2^ of the RDA model including all initial variables. In doing so we prevent inflation of the overall type I error and of the amount of explained variance. After this, we excluded those retained variables with a variance inflation factor (VIF) over 10 (Borcard et al. 2011) to avoid high collinearity. Despite signs of low orthogonality observable among variables in the partial RDA (especially among BIO18, TREE and NDVI, see Results) we chose not to exclude more variables or to apply dimension-reduction treatments like PCA to the environmental space of variables so as to assess their specific and relative contributions to differentiation patterns (McCormack et al. 2010) and discuss signals of adaptation with higher confidence. The final selected ecological variables were used as explanatory variables in two RDAs, with and without subtracting the effects of neutral processes, which were approximated by the first two principal components of the PCA of population structure based on selectively neutral loci (see above). As response variables, we used the same SNP dataset used for the PCA, but excluding LD and neutrality filters. SNP data were coded as counts of the alternative allele for each position (*i.e*., 0, 1 or 2 copies) with VCFTOOLS and transformed following Patterson et al. (2006). Statistical significance was obtained using a permutation-based procedure with 10,000 permutations, assuming α = 0.01. We also used variance partitioning as implemented in the vegan R-package to estimate (i) the total proportion of genomic variation explained by ecological variables alone; (ii) by neutral structure alone; and (iii) the effects of both sets of variables. Finally, we repeated the whole RDA treatment for a subset of 87 SNPs identified as selectively divergent by BayeScan, with no conditional treatment. The analyses were conducted in R version 3.2.2 (code included in Appendix I, Supplementary Material).

**Table 3.**
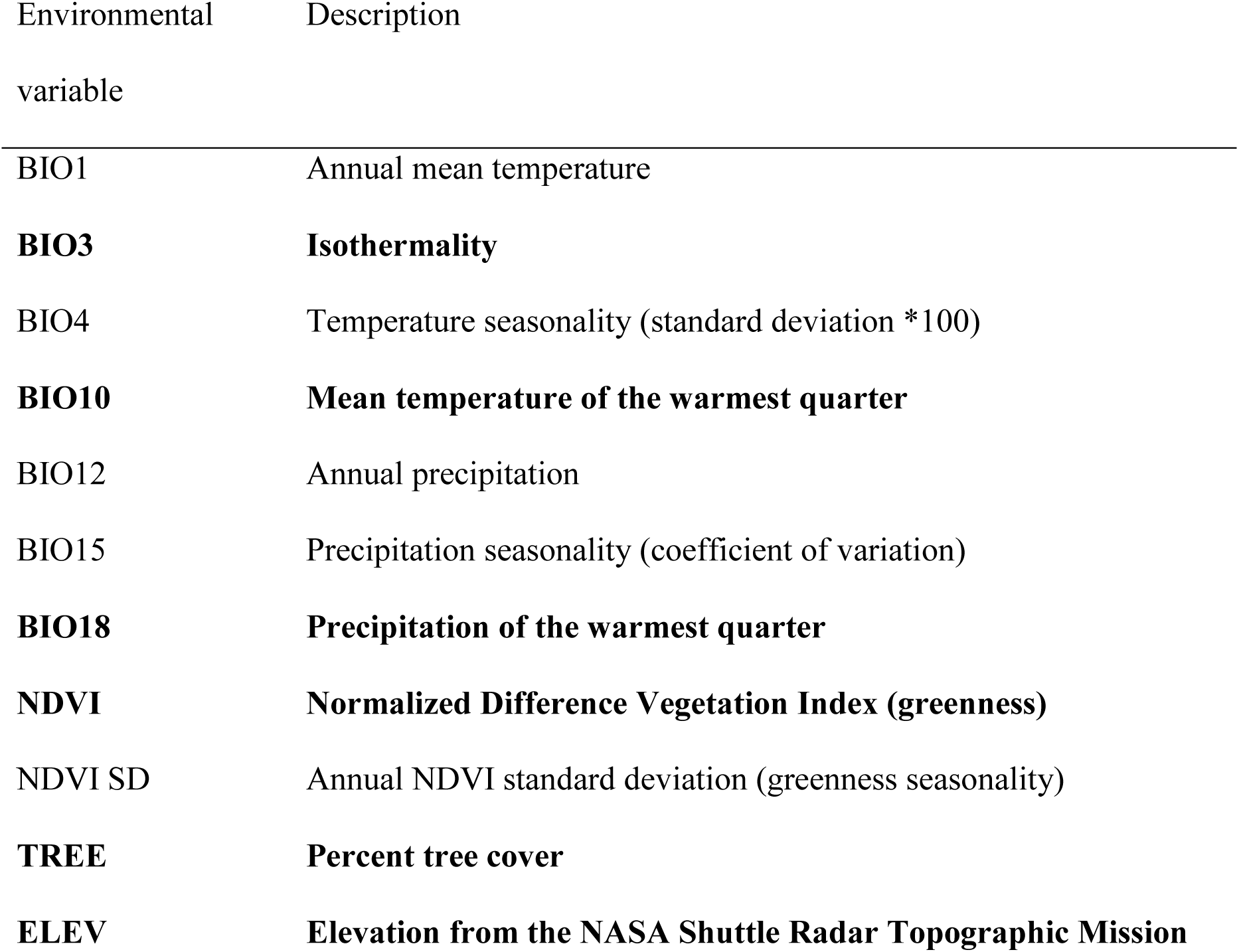
Set of environmental variables included in the initial stepforward selection method. Significant variables retained by the method are shown in bold.

| Environmental variable | Description |
| --- | --- |
| BIO1 | Annual mean temperature |
| **BIO3** | **Isothermality** |
| BIO4 | Temperature seasonality (standard deviation *100) |
| **BIO10** | **Mean temperature of the warmest quarter** |
| BIO12 | Annual precipitation |
| BIO15 | Precipitation seasonality (coefficient of variation) |
| **BIO18** | **Precipitation of the warmest quarter** |
| **NDVI** | **Normalized Difference Vegetation Index (greenness)** |
| NDVI SD | Annual NDVI standard deviation (greenness seasonality) |
| **TREE** | **Percent tree cover** |
| **ELEV** | **Elevation from the NASA Shuttle Radar Topographic Mission** |

### Niche divergence tests

To further explore patterns of ecological divergence in the Oregon Junco, we tested for niche divergence applying the method developed by McCormack et al. (2010), a method that allows us to examine each environmental variable separately. To avoid a loss of statistical power due to multiple analyses, we conducted three specific comparisons of forms presenting different patterns of genetic divergence and geographical settings, including: (i) *townsendi* and southern *thurberi* forms, in order to estimate niche divergence between geographically isolated, genetically differentiated forms; (ii) the ecologically divergent *pinosus* with the parapatric northern *thurberi* form, to further test a possible case of isolation-by-adaptation; and (iii) northern and southern populations of *thurberi*, as conspecific extremes of a potential adaptive gradient. We used occurrence points from our own georreferenced field sampling records, and this set of occurrence records was further revised to avoid spatial autocorrelation and to match the spatial resolution of environmental variables (1-km grid). Our final dataset comprised 80 localities: *pinosus* (n = 14), *thurberi* north (n = 26), *thurberi* south (n = 19), and *townsendi* (n = 21). We decided to improve quality (geographic accuracy) vs. quantity (number of occurrence records), by using fewer data but with higher spatial accuracy (Engler et al. 2004). To generate a background dataset for each population, we drew 1000 random points from a background representing the geographic range of each junco population. In order to select an appropriately sized area for the niche divergence tests, we included accessible habitats according to the dispersal ability of each population (Soberon and Peterson 2005). We generated background samples from a 100-km “buffer zone” around known occurrences (Warren et al. 2008). For populations with small ranges or small dispersal ability (*thurberi* south, *pontilis* and *townsendi*) we restricted the buffer zone to 10 km to reduce spatial inaccuracies in the null distribution (Barve et al. 2011), after testing different buffer sizes to test the robustness in delimiting accessible areas for juncos. Next, we extracted the environmental data (same as the data used for the RDA, see above) for both occurrence points and random background points from within the geographic range of each junco form. Niche divergence and conservatism were tested by comparing the observed environmental differences among forms against a null model of background divergence (generated by calculating the difference between background points using a bootstrapping approach and 1000 resamples) for each environmental variable using a two-tailed test. We conducted all the analyses in R 3.2.2.

### Genome scans

We performed genome scans for different Oregon junco forms using BayeScan. In order to obtain the chromosomic positions of the SNPs, we mapped the GBS reads against the zebra finch (*Taeniopygia guttata*) genome v87 available in Ensembl (Yates et al. 2016), applying the same set of tools and parameters as for mapping against the junco genome. Using the same set of samples as in the PCA and the RDA, we conducted the analysis for all forms together; for *townsendi* against *pontilis*; and for *townsendi* against all *thurberi* (see Table 2 for final dataset sizes). For each of these matrices, we retained only biallelic SNPs with coverage between 2 and 100 and a genotyping phred quality score over 40. Positions with less than 25% of the individuals genotyped were removed from each data matrix, along with those presenting a MAF below 0.05. Once again, we implemented a p-value threshold for HWE of 10^−4^ to filter out false variants arisen by the alignment of paralogous loci. We ran BayeScan with the same settings used for filtering out SNPs under selection in population structure analyses, but implemented a more conservative 10% FDR for outlier detection. Genome scan plots were conducted in R 3.2.2 using the package qqman (Turner 2014).

## Results

### Neutral genetic structure

The plot of the first two principal components from the PCA revealed four distinct clusters in the Oregon junco group. The most differentiated groups were *townsendi* and *pontilis* from Baja California, which formed two highly divergent clusters with respect to each other and to other populations. A third, highly differentiated group corresponded to *pinosus* from coastal California, showing less differentiation than the Baja California forms with respect to a fourth cluster, which included all the remaining forms in the PCA (Fig. 1B). Within this fourth cluster, southern *thurberi* individuals presented certain degree of differentiation from the rest of the forms, a pattern more conspicuous when plotting the third and fourth components, which also revealed a slight signal of divergence in the *oreganus* form (Fig. S1 in Supplementary Information).

The STRUCTURE results were generally congruent with the PCA. The K = 2 plot recovered *townsendi* as an independent population, which shared a considerable amount of variance with *pontilis*. In the analysis for K = 3, *pontilis* separated, and K = 4 identified the same four main clusters seen in the PC1 vs. PC2 plot (Fig. 1B), revealing *pinosus* as an independent genetic cluster. The plot for K = 5 clearly captured the differentiation of the southern *thurberi* form in a fifth cluster, and in K=6, *oreganus* appears as an independent northern group with all individuals from northern *thurberi* and especially *montanus* and *shufeldti* forms showing an increasing probability of assignment to the *oreganus* cluster from south to north (Fig. 2).

Patterns of divergence in Nei’s distances and *F_ST_* values among forms were highly congruent with previous results, with *townsendi*, *pontilis* and *pinosus* showing the highest values for both indices, and northern forms showing lower levels of differentiation, especially between southern and northern *thurberi* individuals (Table S1).

### Forward selection of explanatory variables

Out of eleven potentially relevant ecological variables for juncos, six were retained after the forward selection method intended for excluding non-significant effects. Retained variables included isothermality (BIO3), mean temperature of the warmest quarter (BIO10), mean precipitation of the warmest quarter (BIO18), vegetation cover (TREE), greenness (NDVI), and elevation (ELEV) (Table 3). None of these variables was excluded due to excessive correlation as VIF values were below 10 (maximum recovered VIF = 5.75).

### Redundancy analysis and variance partition

All six ecological variables retained in the forward selection method were included as explanatory variables in the RDA and the partial RDA models. RDA computes, in successive order, a series of axes that are linear combinations of the explanatory variables, and that best explain the variation in the matrix of response variables (Borcard et al. 2011). Six RDA axes (named RDA1 to RDA6 hereafter, ordered by the amount of variance explained by each one, as reflected by the adjusted R^2^) explained 6.26% of the total genetic variance in the non-conditioned model, and 1.18% when removing the effects of neutral genetic structure (Table 4). The amount of explained variance increased to 36.61% when using only BayeScan outliers as response variables (Table 4, Table S2). The permutation tests for the RDA models yielded a p-value below 0.001 in all three analyses, confirming the significance of the constraining variable effects.

**Table 4.** RDA loads of the constraining variables in the first two axes and their explained variance for each one of the RDA models. The total variance explained by the full model (adjusted R^2^ for the resultant six axes) is shown for each analysis. In all of the three analyses, the p-value for the full models was below 0.001. See Table 3 for variable definitions.

|  | Non conditioned RDA |  | Partial RDA |  | RDA of BayeScan outliers |  |
| --- | --- | --- | --- | --- | --- | --- |
| **Total explained variance (adjusted $R^2$)** | 6.26% |  | 1.18% |  | 36.61% |  |
| **Variable** | RDA1 | RDA2 | RDA1 | RDA2 | RDA1 | RDA2 |
| BIO3 | 0.124 | 0.239 | 0.839 | -0.427 | 0.214 | 0.763 |
| BIO10 | 0.108 | -0.610 | 0.359 | -0.679 | 0.221 | 0.374 |
| BIO18 | 0.052 | 0.021 | -0.452 | 0.501 | -0.031 | -0.994 |
| ELEV | 0.646 | -0.031 | -0.324 | -0.301 | 0.629 | 0.420 |
| NDVI | -0.641 | 0.588 | -0.320 | 0.199 | -0.731 | -0.277 |
| TREE | -0.710 | 0.320 | -0.270 | 0.326 | -0.776 | -0.423 |
| **Variance** |  |  |  |  |  |  |
| Eigenvalue | 1304.262 | 567.705 | 481.320 | 350.714 | 63.655 | 12.657 |
| Proportion explained | 0.025 | 0.011 | 0.003 | 0.002 | 0.250 | 0.050 |
| Cumulative proportion | 0.025 | 0.037 | 0.003 | 0.005 | 0.250 | 0.300 |

Loadings of ecological explanatory variables on each of the axes varied across the three different RDA models (Fig 4, Table 4, Table S2). In the non-conditioned RDA, the RDA1 axis had a large negative contribution of TREE and NDVI, and loaded positively on elevation (Fig. 4A). The RDA2 axis loaded mostly on BIO3 and BIO10. The plot of per-individual projections on these two axes revealed a pattern generally similar to the PCA. The forms *townsendi*, *pontilis* and to a lesser extent *pinosus*, showed distinctive high values of correlation with both axes. The remaining forms showed similar correlation patterns with respect to RDA1, with southern *thurberi* individuals showing a clear association with RDA2 (Fig. 4A).

**Figure 4.**
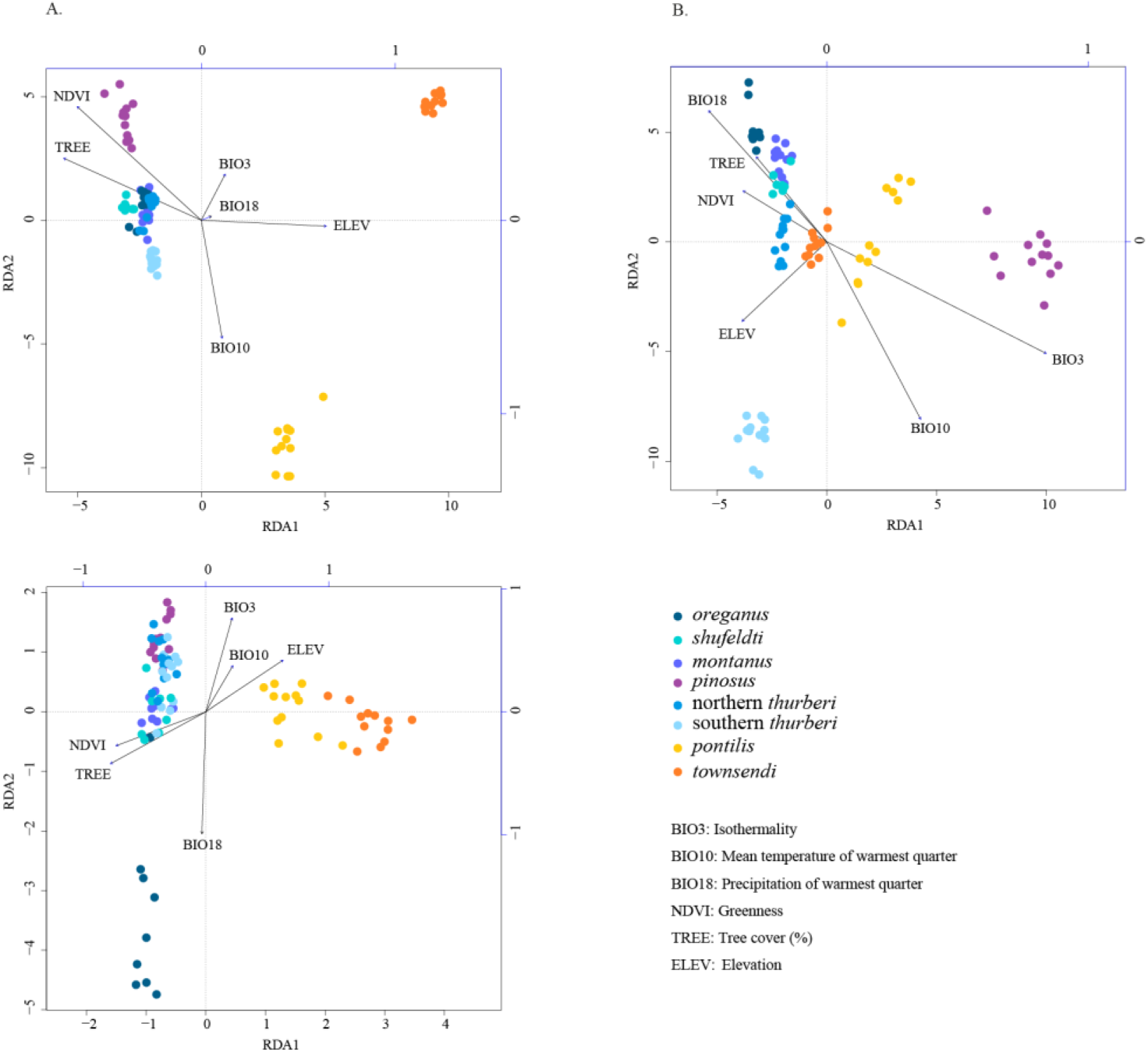
Genetic-environment association analyses in the Oregon junco. Points represent the projection of individual genotypes on the first two RDA axes. Marker colors correspond to those on the range map on Fig. 2A. The explanatory variables are shown within the space defined by RDA1 and RDA2 by labeled vectors. Their contribution to each axis is represented by the length of their orthogonal projections over the scale bars along top and right sides of the graphs. Arrows indicate the direction of the gradient of variation for the corresponding environmental parameter. The value for each sample point on each explanatory variable can be obtained by an orthogonal projection on the corresponding plotted vector. (A) First two RDA axes of a non-conditioned RDA based on 11,261 SNPs. (B) First two RDA axes of a partial RDA based on 11,261 SNPs conditioned by neutral genetic structure, approximated by the first two PCs of a PCA based on neutral markers. (C) First two RDA axes of a non-conditioned RDA based on 87 SNP outliers identified in a BayeScan analysis.

In the partial RDA, RDA1 had a large contribution of BIO3, while RDA2 loaded mostly on BIO10 and to a lesser extent, BIO18 and TREE (Fig. 4B). Plotting these first two RDA axes revealed patterns of genetic correlation especially related to the first RDA axis for *pinosus*, which consequently presented the strongest association with isothermality. The individuals from the southern population of *thurberi* showed a more pronounced negative association with RDA2 than in the non-conditioned RDA, further evidence of a positive correlation with mean temperature of the warmest quarter. Northern forms of the Oregon junco (northern *thurberi*, *shufeldti*, *montanus* and *oreganus*) also separated along the second RDA axis, with the northernmost *oreganus* showing the strongest association with the particularly conspicuous mean precipitation of the warmest quarter gradient, a pattern that was not visible in the non-conditioned RDA. *Townsendi* from Baja California occupied positions closer to the origin of coordinates, suggesting a lower association between environmental and genetic variance (Fig. 4B).

In the RDA based on outlier loci potentially under selection, the first axis showed moderate negative contributions from TREE and NDVI, and also a positive contribution of ELEV (Fig. 4C). Variance in BIO18 was almost entirely captured by RDA2, which also had a relatively high, negative contribution from BIO3. The plot showed a pattern of correlation between *pontilis* and *townsendi* along RDA1, while genetic variance in *oreganus* appeared strongly associated with the gradient of mean precipitation of the warmest quarter along RDA2. The rest of Oregon junco forms (and one atypical *oreganus* individual) were distributed in an opposite fashion, with small differences in their patterns of correlation with environmental variability captured in the second axis of the RDA (Fig. 4C).

The variance partition analysis showed that climate and neutral structure together explained 7.17% of the total genetic variability (fractions A+B+C, Fig. 5). Since variable sets are not orthogonal, a 5.08% of variation was explained jointly by the environmental data and the first two components of the PCA based on neutral genetic positions (fraction B, Fig. 5). As recovered in the partial RDA, environmental variables alone explained 1.18% of the total variance (fraction A, Fig. 5), while the non-overlapping fraction of neutral genetic structure explained 0.91% of the variability in the SNP dataset (fraction C, Fig. 5). The p-value computed through the 10,000-step permutation test for each individual fraction was below 0.001 in all cases, thus confirming the significant effects of both variable sets.

**Figure 5.**
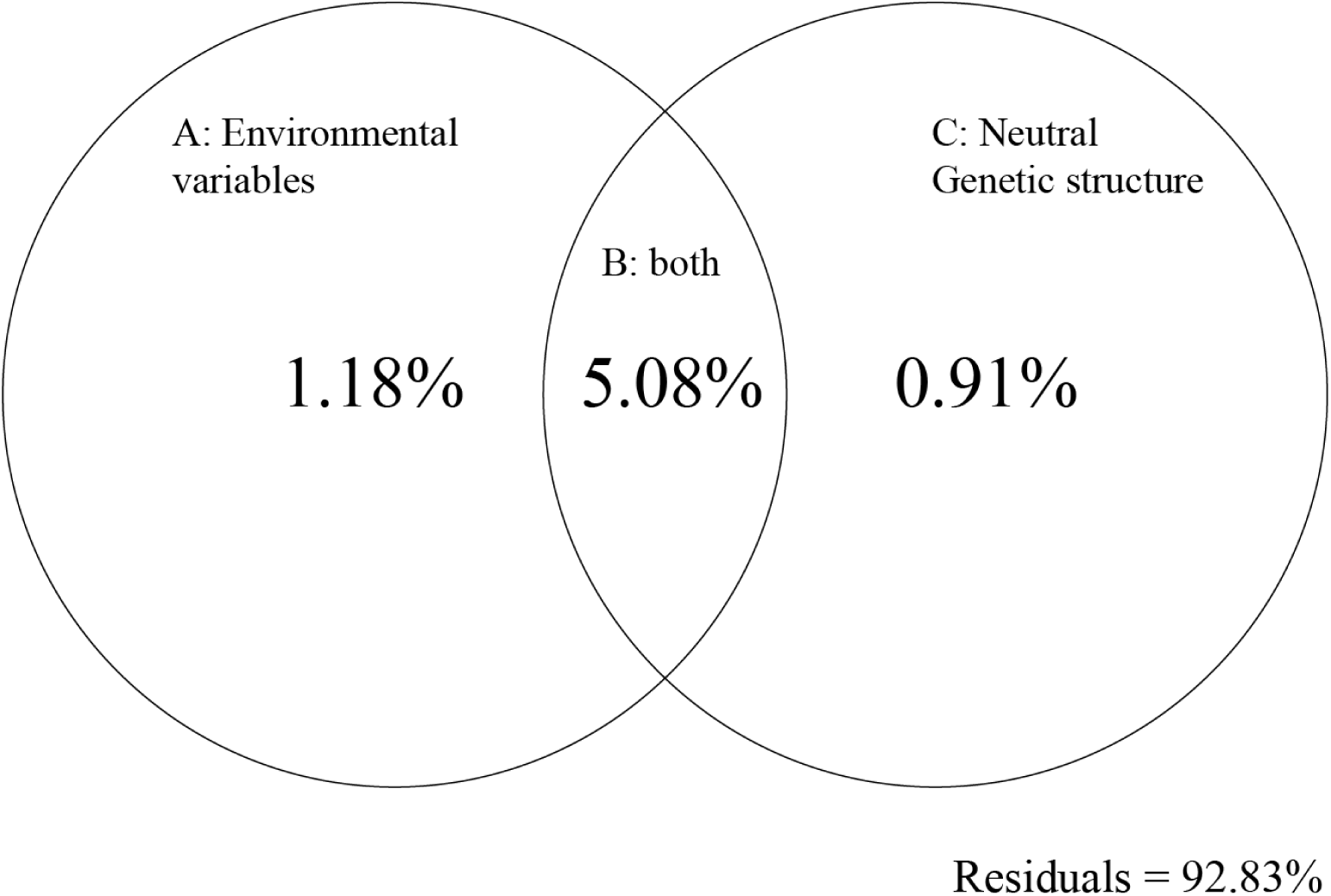
Plot of the fractions of the genetic variability in Oregon samples explained by (A) environmental variables alone; (B) the overlap of both environmental variables and neutral structure; and (C) neutral genetic structure alone; and the unexplained genetic variability (residuals). P-values computed through a 1000-step permutation test for the fractions A, B and C, were below 0.001 in all cases.

### Niche divergence tests

We tested for niche divergence and conservatism on each of the environmental variables. We found significant niche divergence between *pinosus* and northern *thurberi* for three of the six environmental variables analyzed (isothermality, mean precipitation of the warmest quarter and elevation; Table 5). When considering northern *thurberi* vs. southern *thurberi*, we found significant divergence for mean temperature of the warmest quarter and conservatism for isothermality (Table 5). NDVI was the only variable that exhibited significant divergence between *townsendi* and *thurberi* south (Table 5).

**Table 5.** Results from the niche divergence test for *pinosus* vs. northern *thurberi*, northern vs. southern *thurberi* and southern *thurberi* vs. *townsendi*. Variables showing significant divergence (Diverged) or conservatism (Conserved) are shown in bold (p-value < 0.05). e.d.b.b.: expected divergence based on background.

|  | Variable | Result | p-value | Observed mean difference | Background mean difference |
| --- | --- | --- | --- | --- | --- |
| *pinosus* vs. | TREE | e.d.b.b. | 0.636 | 3.95 | 7.17 |
| northern *thurberi* | BIO10 | e.d.b.b. | 0.718 | 20.40 | 17.34 |
|  | **BIO3** | **Diverged** | **0.000** | **16.78** | **9.45** |
|  | **BIO18** | **Diverged** | **0.000** | **41.92** | **27.24** |
|  | **ELEV** | **Diverged** | **0.032** | **1329.81** | **978.26** |
|  | NDVI | e.d.b.b. | 0.550 | 192.67 | 428.18 |
| northern vs. | TREE | e.d.b.b. | 0.580 | 2.11 | 4.52 |
| southern *thurberi* | **BIO10** | **Diverged** | **0.030** | **41.85** | **23.87** |
|  | **BIO3** | **Conserved** | **0.002** | **1.37** | **2.87** |
|  | BIO18 | e.d.b.b. | 0.860 | 15.02 | 14.25 |
|  | ELEV | e.d.b.b. | 0.474 | 202.60 | 140.29 |
|  | NDVI | e.d.b.b. | 0.144 | 258.89 | 1019.97 |
| southern *thurberi* | TREE | e.d.b.b. | 0.536 | 15.01 | 12.25 |
| vs. *townsendi* | BIO10 | e.d.b.b. | 0.412 | 47.88 | 43.03 |
|  | BIO3 | e.d.b.b. | 0.472 | 2.47 | 2.89 |
|  | BIO18 | e.d.b.b. | 0.188 | 74.77 | 67.26 |
|  | ELEV | e.d.b.b. | 0.786 | 750.22 | 717.21 |
|  | **NDVI** | **Diverged** | **0.012** | **1597.62** | **574.28** |

### Genome scans

The BayeScan survey comparing all Oregon junco forms together detected 32 SNPs potentially under divergent selection, and 5 significant SNPs potentially under balancing selection. In the two pairwise comparisons *townsendi* vs. *pontilis* and *townsendi* vs. *thurberi*, 20 and 30 significant SNPs with signs of having diverged under selection were detected, respectively, and no significant SNPs under balancing selection were found in either case. SNPs potentially differentiated under divergent selection appeared distributed across the genome in all comparisons, without obvious signs of heterogeneity among regions (Fig. 6). Chromosomes 1B and 16 harbored no SNPs so they are not shown in the plot.

**Figure 6.**
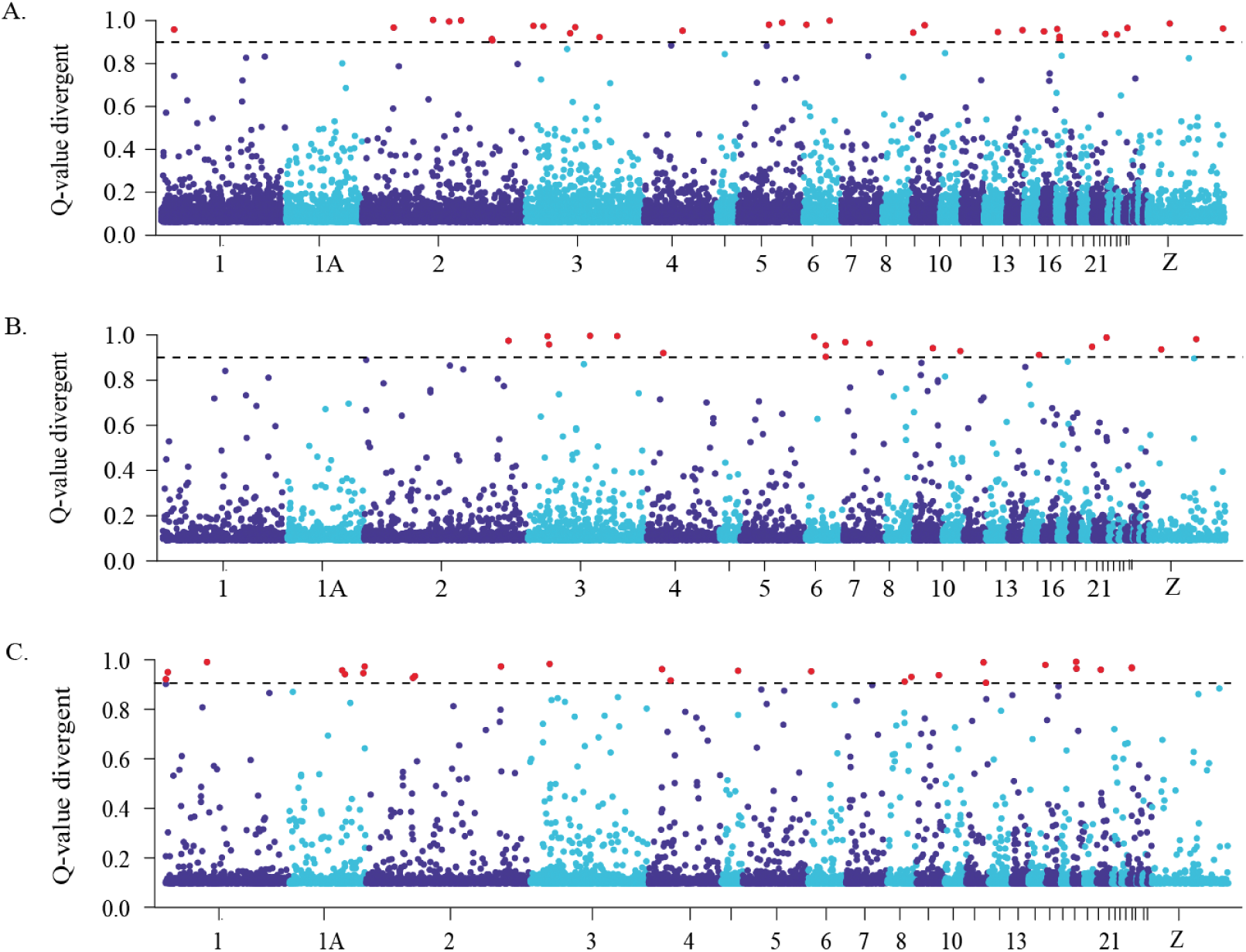
Plot of per-SNP posterior probability of divergence mediated by selection (shown as 1 – Q-value as computed by BayeScan) in (A) all Oregon junco forms together; (B) *townsendi* against *pontilis* and (C) *townsendi* against all *thurberi*. Loci above the dotted line (in red) are those below a false discovery rate of 10%.

## Discussion

### Neutral population structure and local adaptation explains genomic variance among Oregon junco forms

Our results reveal that both neutral and selective factors have played a role in driving divergence among Oregon junco populations, and that the relative contributions of geographic isolation and environment-driven selection are not uniform across the distribution range of the complex. Environmental variables explained 1.18% of genomic variation when controlling for population structure, and environment and neutral structure together accounted for 7.17% of the variability in the 11,256 SNP matrix. The remaining 92.83% of the variance corresponds potentially to loci under balancing selection or selective pressures not represented in our ecological variables, and to neutral variation shared by all Oregon junco forms because of their close relatedness and/or gene flow among them. The amount of variance explained solely by environmental variables in our study was comparable to the values reported in studies applying RDA to detect specific correlations between genomic variation and a given set of potentially correlated variables, as shown in plants (e.g. Lasky et al. 2012; Vincent et al. 2013; De Kort et al. 2014) or other avian species (Safran et al. 2016; Szulkin et al. 2016). Previous studies on birds have used simple spatial variables such as geographic distance to control for the effects of spatial autocorrelation (Safran et al. 2016; Szulkin et al. 2016). Here, we controlled for genome-wide patterns of neutral variation by subtracting the variance captured by the first two PCs of a PCA based on neutral genome-wide SNPs, a method which should better account for population history and structure, including changes in effective population size, geographic isolation and related effects (Forester et al. 2016). Since spatial autocorrelation is usually reflected by neutral genetic structure, we did not include spatial covariates to avoid over-conditioning the model (Rellstab et al. 2015). Given that only a small fraction of the surveyed genome is expected to be related to genes coding for climatic adaptation or linked to them (Meirmans 2015), a significant 1.18% of association between genomic variation and environmental variability in the conditioned (partial) RDA over only 11,261 genome-wide distributed SNPs is a compelling signal of local adaptation.

### Genetic-environment association patterns in the diversification of the Oregon junco

The RDA revealed a number of strikingly different patterns of covariation between genetic variance and ecological variables likely to have played a role in Oregon junco diversification, especially when the effects of population history were removed. The forms *pontilis* and *townsendi* from Baja California, markedly isolated in terms of geography and neutral genetic variability, presented a low genetic-environment association when controlling for population history. This suggests that the differentiation between *townsendi* and *pontilis* is due largely to geographic isolation, in this case caused by unsuitable desert habitat in the lowlands surrounding their respective mountain ranges, a pattern also known as isolation by resistance (IBR, McRae and Beier 2007). In this scenario, our results suggest that differentiation is caused by drift under conditions of small population size and reduced gene flow, rather than divergent selection due to local adaptation, fitting the classic allopatric speciation model (Mayr 1942, 1963; Coyne and Orr 2004). This hypothesis is consistent with the niche divergence test comparing *townsendi* with southern *thurberi*, for which all tested variables but NDVI showed no signal of divergence beyond expectations based on background divergence.

The form *pinosus* showed considerable neutral genetic structure and a conspicuous pattern of genetic-environment association in both non-conditioned and partial RDA. When controlling for population structure, *pinosus* individuals showed high positive correlation values with isothermality, while correlating negatively with elevation. Indeed, *pinosus* presents the highest isothermality values, and the second lowest elevation after *oreganus*, reflecting a tolerance for low elevation conditions that are absent in neighboring *thurberi*, a pattern also recovered in the niche divergence test. Unlike other differentiated forms like *pontilis* and *townsendi*, *pinosus* does not show high geographic isolation, and zones of intergradation with *thurberi* have been described (Miller 1941). Neutral population divergence despite the absence of geographic barriers to gene flow along with signs of local adaptation is a pattern consistent with isolation-by-adaptation, where barriers to gene flow may have arisen as individuals adapted to the distinct habitat of the coastal mountains of California. Niche distinctiveness and the genetic-environment association pattern of this form is thus congruent with a combination of warm latitude, low elevation and coastal influence that has seemingly resulted in the adaptive differentiation of *pinosus* from the rest of the Oregon junco taxa. As a result, differentiation by drift may have led to positive correlations between adaptive and neutral genetic divergence (Nosil et al. 2008).

The southern *thurberi* individuals from Mont Laguna showed high overlap in terms of neutral genetic structure with northern *thurberi* and other boreal forms, and only slight differences in their genetic-environment association patterns when no controls for confounding factors were implemented. The Mount Laguna site represents the southernmost tip of the *thurberi* range in Southern California, which extends northward and reaches Oregon, forming a relatively continuous distribution (Miller 1941; Nolan et al. 2002), suggesting potentially high gene flow. However, the partial RDA revealed a distinctive pattern of high correlation with the mean temperature of the warmest quarter for the southern *thurberi* juncos, differentiating them from the rest of Oregon forms. They also correlated negatively with the mean precipitation of the warmest quarter. This pattern seems congruent with the habitat of Mount Laguna, and in general with the southern inland range of Oregon juncos, quite arid during summer but subject to snowfalls in winter due to the high elevations (Miller 1941), contrasting sharply with the more climatically moderate coastal and northern populations. The limited neutral genetic structure between *thurberi* range extremes but considerable differentiation in the genetic-environment association patterns is consistent with a process of local adaptation across a selective gradient (Forester et al. 2016), in which selection is the prominent evolutionary force driving differentiation (Haldane 1948; Slatkin 1973; Nagylaki 1975; Felsenstein 1976) while neutral alleles may move freely across space. The niche divergence test comparing southern and northern *thurberi* populations was significant for mean temperature of the warmest quarter.

The boreal Oregon junco forms including *oreganus*, *montanus*, *shufeldti*, and *thurberi* individuals from Tahoe, California, presented a more conspicuous pattern of local adaptation along a shallow selective gradient. These forms showed very low neutral genetic structure or differences in ecological covariances in the non-conditioned RDA, yet showed an increasing signal of association following their latitudinal distribution in the partial RDA. A strong association pattern emerged especially for mean precipitation of the warmest quarter and for correlated environmental variables of tree cover and greenness, matching quite precisely their latitudinal distribution along a gradient of increasing humidity and vegetation cover. The ecology-related differences in genetic variance, consistent with the taxonomic classification of these forms, is especially relevant considering the relative phenotypic similarity of these taxa, and their apparent intergradation (Miller 1941; Nolan et al. 2002).

GEA methods present a number of limitations, including potentially high rates of type I error (see Supplementary Materials for details). Here, rather than detecting specific loci under selection, we aimed to explore how selection and neutral processes shape the variability in Oregon juncos, but the risk of finding false significant associations between genetic variance and ecological parameters persists. To further test the environmental associations revealed in this study, we implemented a highly conservative approach using only BayeScan outliers as response variables in the model. The non-conditioned RDA based on 87 SNP loci identified by BayeScan as potential targets of divergent selection yielded relatively lower resolution than the partial RDA. BayeScan has been shown to produce relatively few false positives, but it is also a conservative approach, the sensitivity of which decreases with selection strength (Narum and Hess 2011). The RDA suggests that BayeScan correctly identified outliers related to low temperatures and high precipitation for *oreganus* samples, a pattern congruent with the habitat and with the outcomes of previous analyses for this form. It also detected highly differentiated positions in *pontilis* and *townsendi* that correlate with RDA1, but in this case associations with specific environmental variables were lower, and the pattern disappears in the RDA based on the entire SNP dataset when correcting for population structure. This may suggest that BayeScan failed to exclude the effects of demographic history, or in turn, that controlling for the genetic variance captured by the PCA was overly conservative. BayeScan was also less successful in detecting adaptive divergence in *pinosus*, and especially in northern Oregon junco forms, where selection may be weaker or have acted during a shorter period. Nevertheless, the outlier SNP dataset explained 36.61% of the total climatic variability, a considerable amount compared with the full SNP data RDA models, indicating a good fit of the retained outliers to the linear regression on the environmental parameters.

### Interactions among environment, geography and demographic processes result in three different modes of divergence within the Oregon junco lineage

The Oregon junco is one of the six phenotypically and genetically differentiated dark-eyed junco taxa evolved during a northward expansion from Central America after the last glacial maximum, c.a. 18,000 years ago (Milá et al. 2007; Friis et al. 2016). Similar postglacial expansions have been reported for many other bird species (Seutin et al. 1995; Milá et al. 2000; 2006; Hansson et al. 2008; Malpica and Ornelas 2014; Alvarez et al. 2015). However, the population structure documented in this study reveals a variety of different spatial, selective and demographic factors not previously documented in other avian taxa. In light of our genetic-environment association analyses, the patterns recovered by the PCA and STRUCTURE analysis reveal at least three different effects of geography and demographic history interacting to varying degrees with selection in the process of Oregon junco diversification. First, the IBR pattern of differentiation presented by *pontilis* and *townsendi* may suggest that these forms are peripheral remnants of an original, broader distribution of the Oregon juncos, thereafter isolated in ‘sky islands’ of Baja California and diverging predominantly by drift. Indeed, in his thorough monograph on the geographic variation in juncos, Alden Miller (1941) had perceptively suggested early on that the habitat of Oregon juncos from Baja California did not seem to account for their phenotypic differentiation from Californian forms, and that their distinctive traits appeared to be predominantly “historical” (p. 306). The spatial configuration and recovered patterns of neutral and adaptive divergence for *pontilis* and *townsendi* fit the first model of neutral divergence of isolated population in approximately similar habitats (scenario A in Fig. 2). Second, the IBA pattern found in *pinosus* suggests that the current area of intergradation corresponds to a secondary contact zone that formed after diverging in relative isolation, maybe linked to the ancient coastal closed-cone pine forest that has allegedly diminished since the Pleistocene (Miller 1941). The mode of divergence between parapatric populations in different habitats (scenario B in Fig. 2) has been only partially fulfilled, since local adaptation seems to have resulted in reduced levels of gene flow, leading to increased neutral genome-wide differentiation (Nosil et al. 2008; Funk et al. 2011; Flaxman et al. 2014). Third, the geographic continuum represented by *thurberi*, *shufeldti*, *montanus* and *oreganus* is also captured in the STRUCTURE analysis by a gradual signal of differentiation following a latitudinal distribution, suggesting that ongoing gene flow may occur among forms. Combined with the signal of increasing environmental association recovered in the partial RDA, these outcomes are consistent with a process of differentiation driven by local adaptation along a selective gradient in the direction of the northward expansion, fulfilling the third hypothesized mode of divergence for these forms of Oregon junco (scenario C in Fig. 2). Interestingly, ecological association approaches have been shown to perform better along clines of selection where demographic expansions align with the gradient of ecological variables, usually related to latitude (Frichot et al. 2015), as is the case in the *Junco* system.

A relevant aspect of the marked population structure found among Oregon junco forms is that it is based on a relatively small subset of genome-wide SNPs randomly sampled from across the genome, representing a genomic fraction not greater than 0.2% of the total of 1.2 Gb. The clear signal of divergence mediated by environmental factors recovered also in the RDA indicates that divergence may have taken place at the level of the entire genome, suggesting the role of multiple selective pressures during local adaptation along the latitudinally broad and heterogeneous distribution of the Oregon juncos. The presence of outliers potentially under positive selection scattered across the genome seems to support this hypothesis of selection-driven genome-wide divergence, rather than widespread drift among isolated populations. Other examples of such patterns of genomic differentiation due to divergent selection at early (e.g. Parchman et al. 2013; Brawand et al. 2014; Egan et al. 2015) and intermediate (e.g. Riesch et al. 2017) stages of speciation have been reported recently, contrasting with proposed models of speciation initiated by divergent selection in a few, localized genes involved in reproductive isolation (e.g. Nadeau et al. 2012; Poelstra et al. 2014).

## Conclusion

Our analyses reveal the role of both local adaptation and demography in driving rapid diversification during the northward recolonization of western North America by the Oregon junco. The combined effects of a demographic expansion along a selective gradient with a heterogeneous landscape of environmental variability have resulted in a striking array of divergence modes within a single lineage, from isolated forms in Baja California that have differentiated largely by drift in isolated ‘sky islands’, to adaptive diversification along selective gradients with no obvious geographic barriers to gene flow. There is also a compelling example of isolation by adaptation in the case of *pinosus*, where ecological barriers to gene flow seem to maintain its divergence with respect to nearby forms. Genome-wide patterns of divergence indicate that Oregon junco diversification has been driven by multiple ecological factors acting on many loci across the genome, and suggests that selection may promote local adaptation in short periods of time, highlighting the role of adaptive divergence in the early stages of the speciation process. Future analyses of dense sequencing and functional gene characterization will be necessary to further identify adaptive changes promoting barriers to gene flow and reveal the genomic architecture of rapid diversification.

## Acknowledgements

We thank Pau Aleixandre, Jonathan Atwell, Elena Berg, Setefilla Buenavista, Steve Burns, Jatziri Calderón, Adrián Gutiérrez, Alfonsina Hernández, Fritz Hertel, Ellen Ketterson, Adán Oliveras de Ita, César Ríos, Sahid Robles, Vicente Rodríguez, Whitney Tsai, Rich Van Buskirk and Alvar Veiga for invaluable help in the field and lab. We are also grateful to Yoann Anselmetti, Clémentine Francois, Rachel Johnston, Etienne Kornobis, Benoit Nabholz, Sergio Nigenda, Jacqueline Robinson, Marjolaine Rouselle and Robert K. Wayne for their assistance with bioinformatic analyses. We also thank Laura Barrios and Luis M. Carrascal for reviewing the statistical analyses and providing orientation. Funding was provided by grant CGL-2011-25866 from the Spanish Ministry of Science and Innovation to BM.

## Data Accessibility

Genomic data will be deposited in Dryad in short.

## Author Contributions

G. Friis and BM designed the study and carried out field sampling; G. Friis, JM, and BF generated and analyzed genomic data; G. Friis, G. Fandos and AZ generated and analyzed environmental data; G. Friis and BM wrote the manuscript with input from all co-authors.

